# A global post-transcriptional regulator directs *Acinetobacter baumannii* carbon preference and gut colonization

**DOI:** 10.64898/2026.11.07.762903

**Authors:** R. Mason Clark, Rosa L. Sava, Kia J.A. Holt, Michael J. Gebhardt, Lauren D. Palmer

**Affiliations:** Department of Microbiology and Immunology, University of Illinois Chicago College of Medicine, Chicago, IL, USA; Department of Microbiology and Immunology, University of Iowa Carver College of Medicine, Iowa City, IA, USA

**Keywords:** *Acinetobacter baumannii*, CsrA, carbon storage regulator, carbon preference, carbon catabolite repression, gut colonization

## Abstract

*Acinetobacter baumannii* is a multidrug-resistant pathogen that colonizes the gut, increasing the risk of subsequent infection and transmission. Recent work showed that *A. baumannii* catabolizes ornithine, a non-preferred carbon source, to colonize the gut. However, the mechanisms of carbon preference were unknown in *A. baumannii*. Here, experimental evolution identified the conserved RNA-binding protein CsrA as a regulator of carbon preference in *A. baumannii*, a previously unreported CsrA function. RNA immunoprecipitation sequencing (RIP-Seq) revealed hundreds of CsrA-bound transcripts*. A. baumannii* CsrA directly bound and repressed expression of the *ast* arginine/ornithine catabolic operons. The *Pseudomonas aeruginosa* CsrA homolog RsmA repressed arginine/ornithine catabolic operon expression, suggesting this CsrA function extends to other bacterial taxa. Finally, CsrA was required for competitive fitness during *A. baumannii* gut colonization. These findings uncover a previously unrecognized role for a global regulator in *A. baumannii* carbon preference and persistence in the gut, a critical reservoir for multidrug-resistant pathogens.

## Introduction

*Acinetobacter baumannii* commonly causes healthcare associated infections and is the second leading cause of global mortality from antimicrobial resistance.^1–3^ Because of limited treatment options, infection prevention is a critical strategy for addressing *A. baumannii* mortality. Growing evidence supports asymptomatic colonization as an important clinical reservoir for multidrug-resistant (MDR) pathogens including *A. baumannii*.^4,5^ While *A. baumannii* can colonize virtually any body site, gut colonization is the most described site of asymptomatic carriage.^6^ Patients asymptomatically colonized with *A. baumannii* are more likely to develop clinical *A. baumannii* infections, contaminate their healthcare environments, and transmit *A. baumannii* to other cohoused patients.^6^ Therefore, understanding *A. baumannii* gut colonization is critical to develop strategies to prevent colonization and infections.

The mechanisms of *A. baumannii* gut colonization are largely unknown. Antibiotic exposure is a major risk factor for *A. baumannii* gut colonization in humans, while mouse models of *A. baumannii* gut colonization generally require antibiotic treatment prior to inoculation.^6–12^ These findings suggest an undisturbed gut microbiota provides colonization resistance to *A. baumannii*. We recently reported that pathogenic *Acinetobacter* evolved to utilize the amino acid ornithine (Orn), which was required to persist in the gut in a post-antibiotics mouse model.^10^ Together, the data suggested that a complex microbiota competed away preferred nutrient sources, such that *A. baumannii* utilized the non-preferred nutrient source ornithine to persist in the gut.^10^ Additional work from our group showed that the transcriptional regulator AstR is required for *A. baumannii* to activate ornithine catabolism in response to microbiota competition and persist in the gut.^13^ Preferred carbon sources such as glutamate inhibited AstR-dependent activation but did not affect AstR-DNA binding.^13^ Therefore, the carbon preference regulatory mechanisms repressing the arginine/ornithine utilization pathway required for *A. baumannii* gut colonization remain unknown.

The regulators of carbon preference (also referred to as carbon catabolite repression) in *A. baumannii* have not been described.^14^ Here, we expand characterization of the previously reported three tiers of *A. baumannii* carbon source preferences.^10^ Experimental evolution identified CsrA, an RNA-binding protein and global regulator, as a carbon catabolite repressor of arginine and ornithine utilization. While CsrA is a global post-transcriptional regulator with pleiotropic effects, it has not been previously reported to regulate carbon preference.^15^ Genome-wide RNA immunoprecipitation sequencing (RIP-Seq) and additional approaches demonstrated that *A. baumannii* CsrA directly binds *ast* transcripts to repress arginine and ornithine catabolism. These pathways are similarly repressed by the CsrA homolog RsmA in *Pseudomonas aeruginosa*, suggesting a common ancestral function for CsrA that was adapted for carbon catabolite repression in *A. baumannii*. Finally, CsrA was required for *A. baumannii* to persistently colonize the mouse gut. Collectively, these data highlight a new role for CsrA in carbon preference and gut colonization of the MDR pathogen *A. baumannii*.

## Results

### *A. baumannii* preferentially uses carbon sources in three tiers

We previously defined three tiers of *A. baumannii* carbon preference for a limited set of carbon sources by diauxic growth curves: tier 1 (Glu, Gln, Asp, His) were utilized before tier 2 (Succinate), and tiers 1 and 2 were utilized before tier 3 (Arg, Orn).^10^ Here, we further characterized *A. baumannii* carbon preferences. *Acinetobacter spp.* generally cannot catabolize sugars but use a wide variety of other carbon sources including amino acids, organic acids, and alcohols.^14^ Diauxic growth curves with wild-type *A. baumannii* 17978 in M9 minimal media compared 27 carbon sources to glutamate, succinate, and arginine as representatives of tier 1, tier 2, and tier 3 (Figure 1a). The amino acids aspartate (Figure 1b) and proline (Figure 1c) showed overlapping growth curves when supplemented to glutamate medium and concentration-dependent lag phases when supplemented to succinate or arginine medium. This suggests that *A. baumannii* uses these amino acids simultaneously with glutamate and preferentially over succinate and arginine, supporting their classification as tier 1 carbon sources.

**Figure 1:**
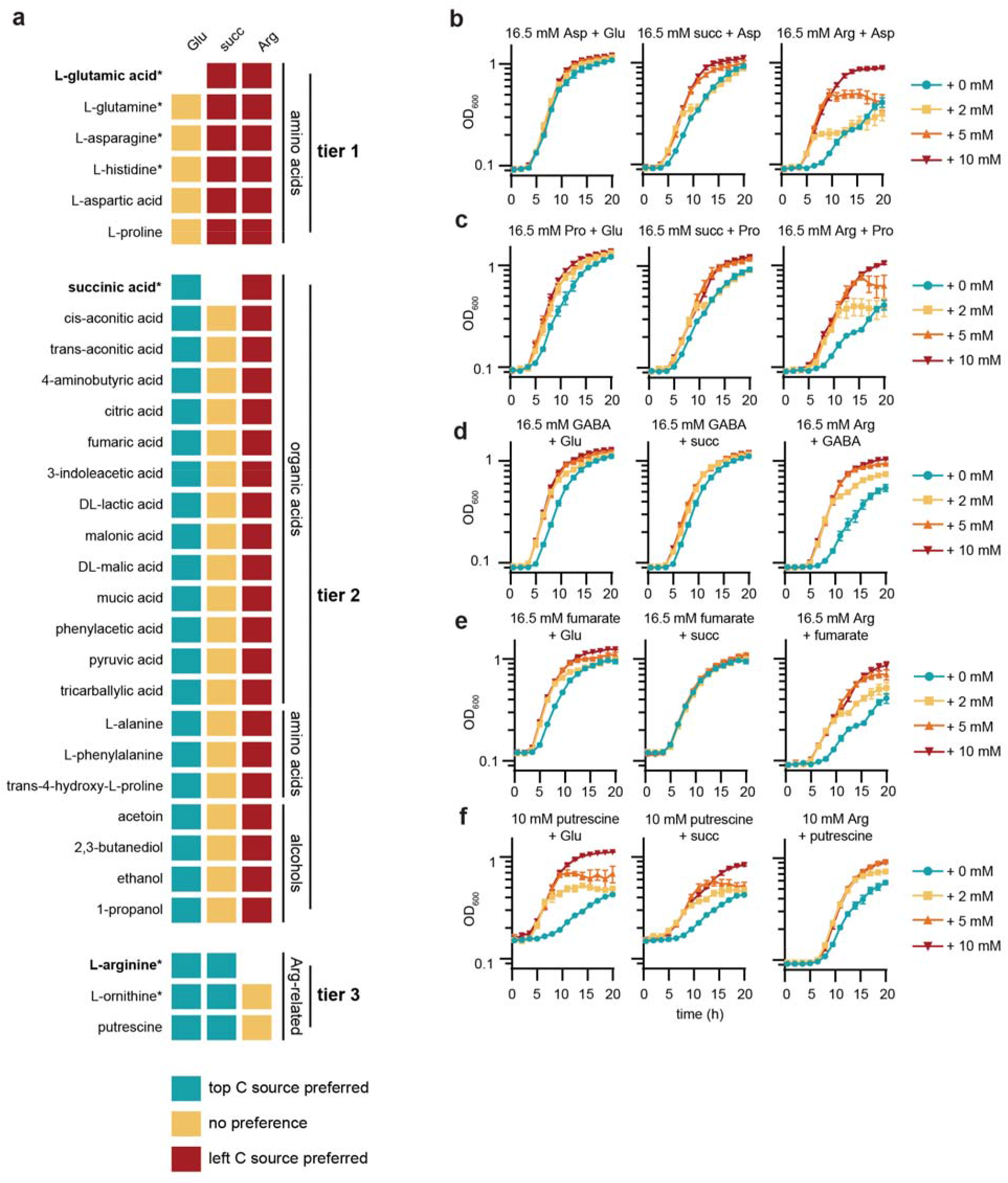
*A. baumannii* displays diauxic growth with carbon sources in three tiers. (a) Summary of tiers of carbon (C) source preference for *A. baumannii* 17978. (b-f) Diauxic growth curves of wild-type *A. baumannii* 17978 in M9 media with the indicated carbon sources. Growth was monitored by optical density at 600 nm (OD_600_, *n =* 3, mean ± SEM, experiments were repeated at least twice with similar results). Succ, succinate; GABA, 4-aminobutyric acid.

Next, the addition of glutamate induced concentration-dependent lag phases in media with excess organic acids 4-aminobutyrate (GABA, Figure 1d), fumarate (Figure 1e), *cis*-aconitate, *trans*-aconitate, citrate, 3-indoleacetate (IAA), DL-lactate, malonate, DL-malic acid, mucic acid, phenylacetic acid (PAA), pyruvate, and tricarballylic acid (Figure 1a, Table S1). These organic acids displayed overlapping growth curves with additions of succinate and induced concentration-dependent lag phases when added to cultures with excess arginine (Figure 1a, Table S1), supporting their assignment to tier 2. Additional tier 2 carbon sources include the amino acids alanine and phenylalanine and the alcohols ethanol, 2,3-butanediol, and acetoin (Figure 1a, Table S1). Thus, the tier 2 carbon sources are largely organic acids with some amino acids and alcohols.

Finally, glutamate and succinate both induced concentration-dependent lag phases when added to cultures with putrescine (Figure 1a, 1f). Putrescine showed overlapping growth curves with additions to cultures with arginine (Figure 1a, 1f). This suggests that putrescine is in tier 3 of *A. baumannii* carbon preference along with arginine and ornithine, all arginine-related compounds. Together, these data support placement of carbon sources into three tiers of preference: tier 1, preferred amino acids, followed by tier 2, mostly organic acids, and finally tier 3, the least preferred arginine-related compounds.

### Experimental evolution revealed CsrA as a regulator of arginine catabolism

Next, we sought to determine the regulator responsible for carbon preference in *A. baumannii*. A targeted approach tested the role of two candidate proteins. First, the catabolite regulatory protein (Crp, also known as cyclic AMP receptor protein) is a global regulator known to contribute to carbon catabolite repression and carbon preference in other species including *Escherichia coli*.^16,17^ While *A. baumannii* Crp has been shown to contribute to natural competence^18^ and is predicted to play a role in biofilm formation,^19,20^ its role in carbon preference was unknown. Second, the catabolite repression control protein (Crc) is known to regulate carbon catabolite repression in related species such as *Acinetobacter baylyi*^21–23^ and *P. aeruginosa*^24^. When grown in minimal media with excess arginine or succinate and limiting glutamate, Δ*crp* and Δ*crc* mutants displayed glutamate-dependent lag phases in growth curves (Figures S1a-b), similar to wild-type. This suggests that neither Crp nor Crc is required for *A. baumannii* carbon preference of glutamate over arginine.

*A. baumannii* 17978 reaches stationary phase relatively slowly in minimal medium with arginine as the sole carbon source (Figure S1c). We hypothesized that slow growth with arginine as the sole carbon source could be due to carbon catabolite repression of the *astGCADBE* operon, the only arginine catabolic pathway encoded by *A. baumannii*. Therefore, we hypothesized that experimentally evolving *A. baumannii* for faster growth with arginine as the sole carbon source may enrich for mutations in genes encoding carbon catabolite repressors. Twelve independent cultures of *A. baumannii* 17978 were subcultured 1:2,000 each day into fresh M9 arginine medium for ten days, totaling approximately 110 generations (Figure 2a). The day 10 populations reached stationary phase in M9 arginine medium faster than the day 1 populations (Figure 2b). Each population was sequenced and analyzed, and results were sorted by frequency to identify loci that had mutations in multiple independent populations.

**Figure 2:**
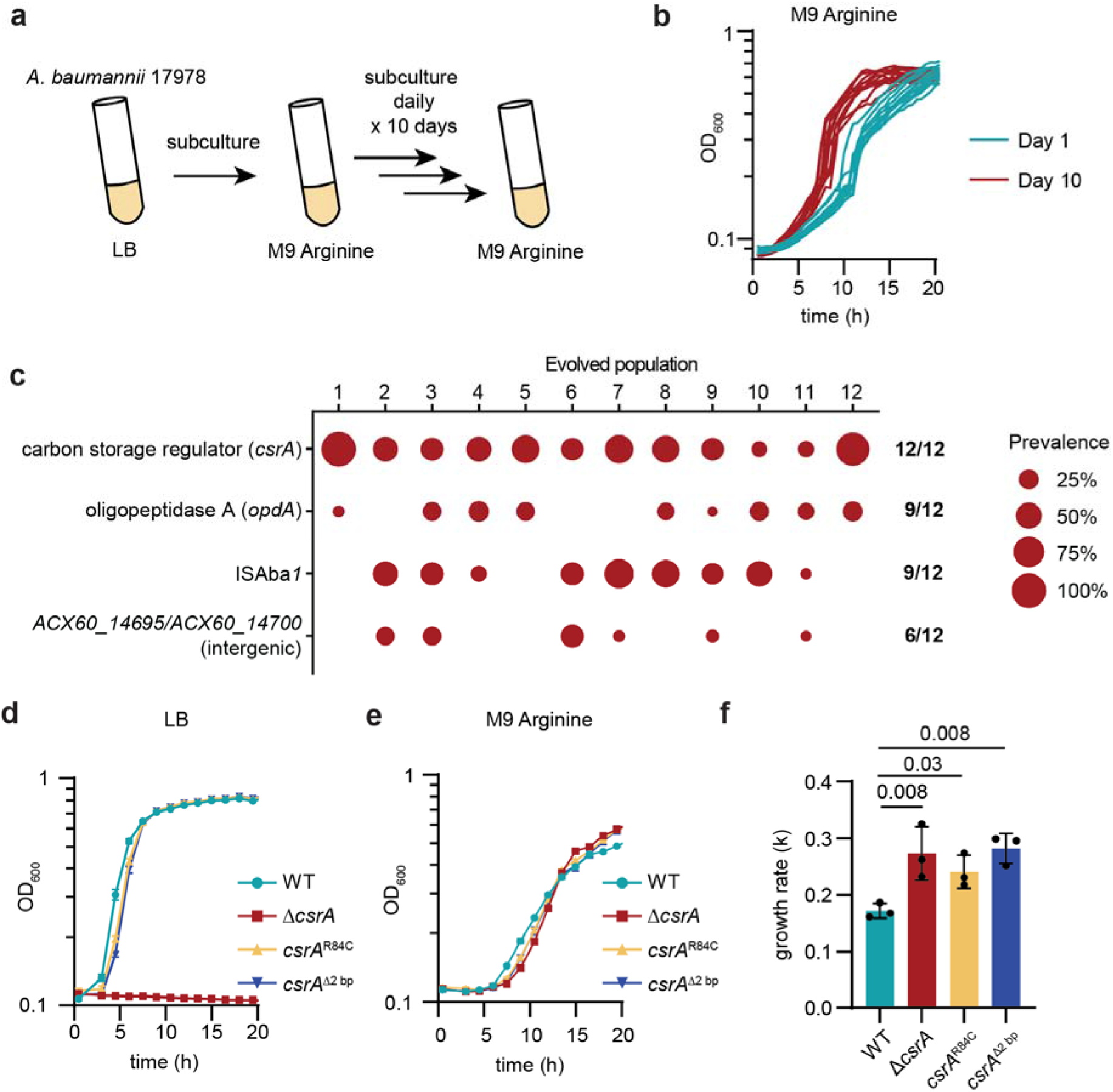
Experimental evolution reveals *csrA* mutations lead to faster growth with arginine as the sole carbon source. (a) Schematic for experimental evolution. (b) Growth of 12 independent populations of *A. baumannii* 17978 in M9 medium with 16.5 mM arginine as the sole carbon source on day 1 and day 10 of the experimental evolution. Each line represents a single population. Growth was monitored by optical density at 600 nm (OD_600_). (c) Genetic loci that were mutated during the experimental evolution. Data were filtered to show loci that have mutations detected in at least six independent populations and prevalence of at least 25% in one population. The size of each dot represents the prevalence of the most prevalent mutation in each locus and population. Fractions indicate the number of independent populations with mutations detected in each locus. (d-e) Growth of wild-type, Δ*csrA*, and reconstructed *csrA*^R84C^ and *csrA*^Δ2bp^ *A. baumannii* 17978 in lysogeny broth (LB) or M9 medium with 16.5 mM arginine as the sole carbon source. Growth was monitored by optical density at 600 nm (OD_600_, *n =* 3, mean ± SEM, experiments were repeated at least three times with similar results). (f) Growth rates (*k*) of *A. baumannii* 17978 strains in M9 medium with 16.5 mM arginine as the sole carbon source shown in Figure 2e (n = 3, mean ± SD, *P* by one-way ANOVA with Holm-Šídák’s multiple comparisons test).

All 12 independent populations evolved mutations in the promoter or coding sequence of carbon storage regulator A (*csrA*) (Figure 2c). The second most common mutation was in oligopeptidase A (*opdA*) which was found in 9/12 populations. An in-frame deletion mutant of *opdA* (Δ*opdA::*Kn) grew similarly to wild-type *A. baumannii* in M9 arginine medium (Figure S1d), and thus *opdA* was not further analyzed. The other two genes mutated in 6 or more populations were those that make up the mobile genetic element ISAba*1* and in an intergenic region where ISAba*1* transposed in 6 populations. No evolved mutations were found in the genes either upstream or downstream of the site where ISAba*1* transposed, suggesting that this intergenic region may be a high-frequency site for ISAba*1* translocation unrelated to growth with arginine as the sole carbon source. Thus, this locus was not further analyzed.

Given the presence of mutations in *csrA* in every population evolved in M9 arginine media, we hypothesized that CsrA represses arginine catabolic genes in *A. baumannii*. To test this, we created an in-frame deletion-insertion strain (Δ*csrA*::Kn; hereafter Δ*csrA*). Additionally, two *csrA* mutations that occurred in multiple independent populations were reconstructed in a wild-type background to generate isogenic strains (*csrA*^R84C^ and *csrA*^Δ2bp^). As previously described, the Δ*csrA* mutant strain was unable to grow in rich media including lysogeny broth (LB, Figure 2d).^25–27^ However, both reconstructed evolved mutant strains were able to grow in LB, suggesting that these alleles are not complete loss-of-function mutations (Figure 2d). All three *csrA* mutant strains had slight but statistically significantly increased growth rates in M9 arginine media compared to wild-type (Figure 2e-f). These data show that disruption of *csrA* leads to increased *A. baumannii* 17978 growth rates when utilizing arginine as the sole carbon source.

### CsrA is required for arginine carbon catabolite repression in *A. baumannii*

Since CsrA restricts *A. baumannii* growth with arginine as a sole carbon source, we predicted that CsrA mediates carbon catabolite repression of arginine when preferred carbon sources are present. As shown above, wild-type *A. baumannii* 17978 displayed diauxic growth and a glutamate concentration-dependent lag phase with excess arginine (Figure 3a). However, the *A. baumannii* 17978 Δ*csrA* strain did not display a glutamate concentration-dependent lag phase with excess arginine, ornithine, or putrescine (Figure 3a-c). CsrA was also required for the glutamate concentration-dependent lag phase with excess arginine in another historical *A. baumannii* isolate, ATCC 17961 (Figure 3d) and the more recent clinical isolates AB5075^28^ (Figure 3e) and Ab09-003^29^ (Figure 3f). These data show that CsrA is required for the preference of glutamate over tier 3 arginine-related compounds in multiple *A. baumannii* strain backgrounds.

**Figure 3:**
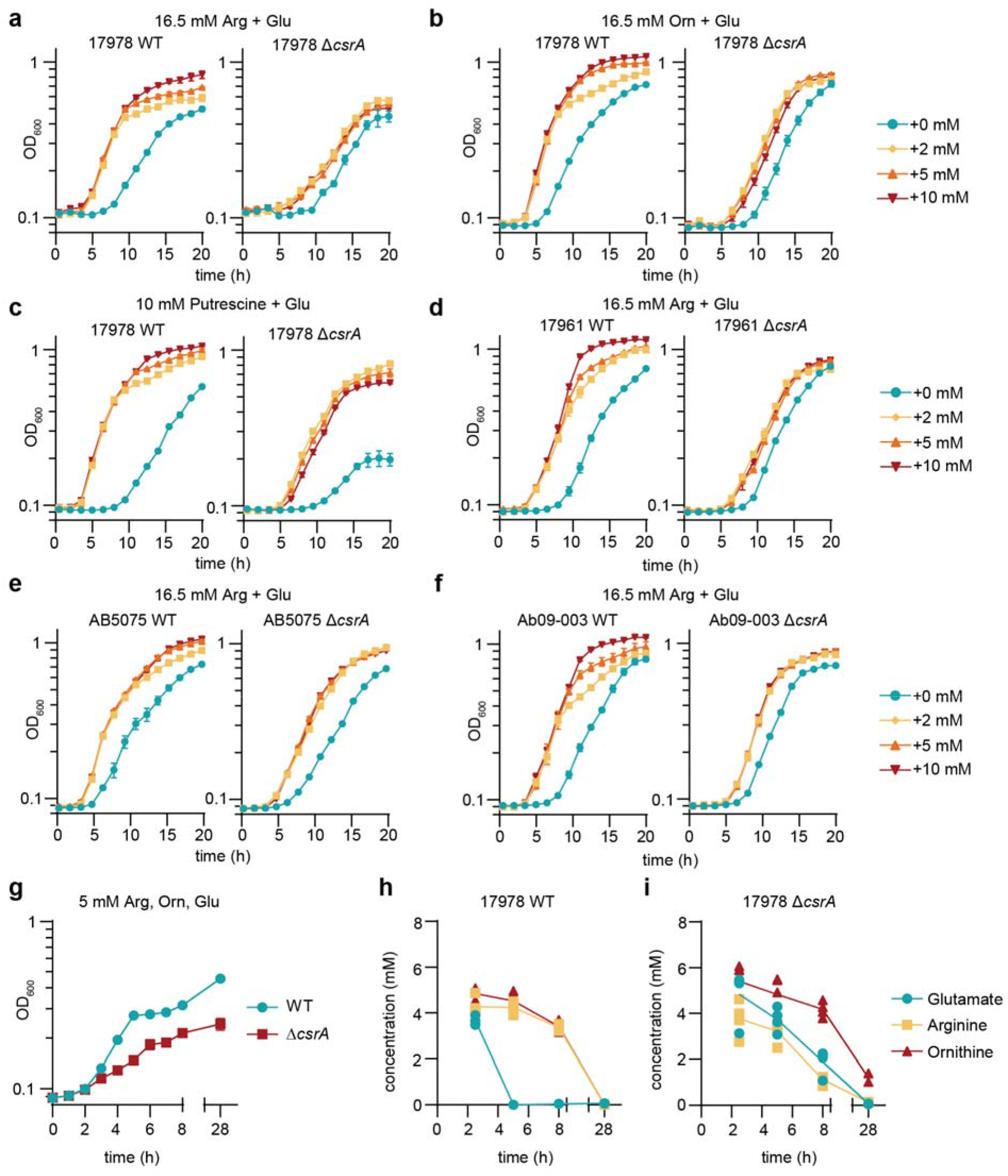
CsrA is required for preference of carbon sources over arginine and ornithine. (a-c) Diauxic growth curves of wild-type and Δ*csrA A. baumannii* 17978 in M9 media with 16.5 mM arginine, 16.5 mM ornithine, or 10 mM putrescine and the indicated concentration of glutamate as the sole carbon source(s). Growth was monitored by optical density at 600 nm (OD_600,_ *n =* 3, mean ± SEM, experiments were repeated at least four times with similar results). (d-f) Diauxic growth curves of wild-type and Δ*csrA A. baumannii* AB5075, ATCC 17961, or Ab09-003 in M9 media with 16.5 mM arginine and the indicated concentration of glutamate as the sole carbon source(s). Growth was monitored by OD_600_ (*n =* 3, mean ± SEM, experiments were repeated at least twice with similar results). (g) Growth of *A. baumannii* 17978 WT and Δ*csrA* in M9 media with 5 mM glutamate, 5 mM arginine, and 5 mM ornithine as the sole carbon sources. Growth was monitored by OD_600_ (*n =* 4, mean ± SEM). (h-i) Concentration of glutamate, arginine, and ornithine in supernatant at 2.5, 5, 8, and 28 hours of growth of *A. baumannii* 17978 WT or Δ*csrA* as shown in Figure 3g (*n =* 4, line represents median).

Next, we sought to characterize if carbon preference relationships outside of tier 3 require CsrA. With limited glutamate (tier 1) added to an excess of succinate (tier 2), *A. baumannii* 17978 diauxie appeared similar between the wild-type and Δ*csrA* strains (Figure S2a). However, *A. baumannii* AB5075 behaved differently: the wild-type strain showed the expected diauxie, but the Δ*csrA* strain did not display any glutamate concentration-dependent lag phase (Figure S2b). This loss of diauxie was also observed with glutamate additions to other tier 2 carbon sources including citrate (Figure S2c), and fumarate (Figure S2d). This suggests that CsrA is not required for glutamate preference over succinate in *A. baumannii* 17978 but is required in AB5075. These experiments indicate that CsrA-mediated regulation of carbon preference between tier 1 and tier 2 varies among *A. baumannii* strain backgrounds.

To test whether the diauxic growth patterns reflect *A. baumannii* carbon preference, we performed targeted metabolomics of wild-type and Δ*csrA A. baumannii* 17978 grown in M9 medium with 5 mM arginine, 5 mM ornithine, and 5 mM glutamate as the sole carbon sources. The wild-type culture displayed diauxie with a lag induced around 5 hours of growth, whereas Δ*csrA* did not display a lag phase (Figure 3g). Amino acid quantification showed that for the wild-type culture, glutamate was depleted by 5 hours of growth, and the arginine and ornithine levels remained near 5 mM until 5 h and began decreasing by 8 h (Figure 3h). This glutamate depletion corresponded to the timing of the lag during the growth of the wild-type culture around 5 hours. However, the Δ*csrA* culture showed glutamate, arginine, and ornithine levels decreasing together throughout the experiment, consistent with a loss of carbon preference (Figure 3i). Together, these data show that CsrA is required for preference of glutamate over arginine and ornithine in *A. baumannii*.

### CsrA directly binds *ast* transcripts in *A. baumannii*

To identify the genes CsrA directly binds in *A. baumannii*, we performed RNA immunoprecipitation and sequencing (RIP-Seq) in *A. baumannii* AB5075 using an VSV-G epitope tagged allele of CsrA (CsrA-V, Figure 4a).^25^ CsrA-V complemented growth of AB5075 Δ*csrA* in LB, suggesting the epitope tag does not disrupt CsrA function (Figure S3a). Enrichment analysis with a 1.5-fold cutoff revealed CsrA associates with 289 RNAs during exponential growth in LB (Figure 4b). Of these targets, 91 are related to metabolism, with regions of enrichment found over *astGCADBE*, *astNOP,* and *astR* (Figure 4c-d, Figure S3b-d), indicating that the involvement of CsrA in carbon preference may be mediated through post-transcriptional regulation of the *ast* loci. Additional targets identified include genes involved in catabolism of tier 1 carbon source histidine (*hutH, hutU*) and tier 2 carbon sources PAA (*paaF, paaJ*) and alanine (*cycA2, dadA, dadX*). These data support CsrA as a global post-transcriptional regulator of processes including carbon catabolism in *A. baumannii*.

**Figure 4:**
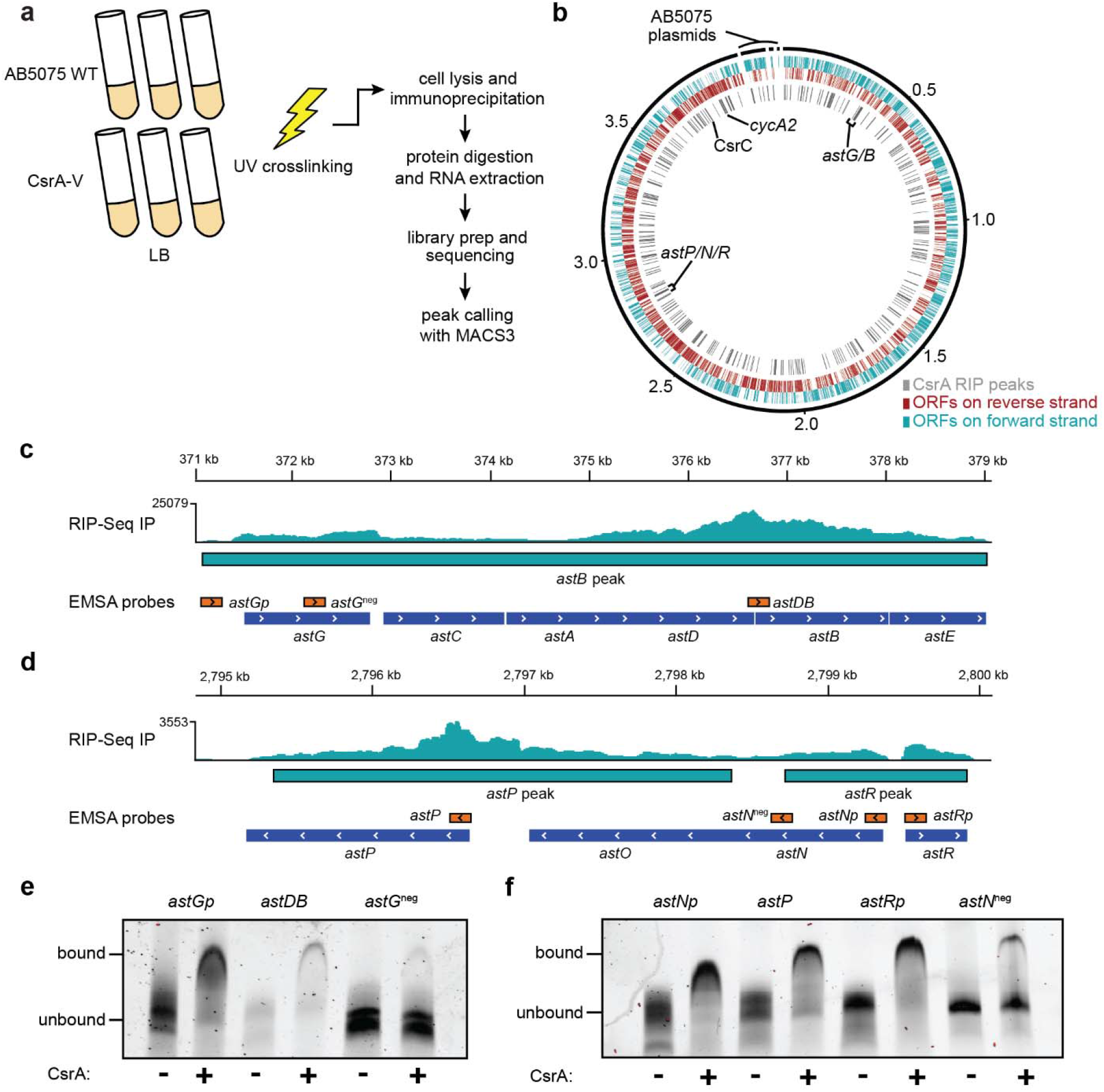
CsrA directly binds *A. baumannii ast* transcripts. (a) RIP-Seq experimental design (b) Circos plot showing open reading frames and RIP peaks with an enrichment of 1.5-fold or greater overlayed on the AB5075 genome. (c-d) RIP-Seq traces showing enriched reads overlayed on the *ast* operons. Teal bars below the read density traces represent significantly enriched transcripts. RNA probes used in RNA EMSAs and the *ast* operon structure are shown below the predicted targets. (e-f) EMSA analysis of purified CsrA-His binding to RNA probes generated from the *ast* transcripts.

The direct binding of *ast* transcripts to CsrA was next validated by RNA electromobility shift assays (EMSA) using purified CsrA with a 6xHis epitope tag appended to the C-terminus (CsrA-His). First, the *A. baumannii* 17978 Δ*csrA* mutant complemented with CsrA-His rescued growth in LB, demonstrating CsrA-His retained function (Figure S4a). EMSA assays showed CsrA-His shifted migration of RNA probes designed from the *ast* transcripts and minimally shifted migration of negative control probes corresponding to regions of each operon with low enrichment by RIP-Seq (Figure 4e-f, Figure S4b-g). Thus, CsrA directly binds RNA at both *ast* loci in *A. baumannii*.

### CsrA represses expression of *ast* transcripts in *A. baumannii* and *P. aeruginosa*

We next assessed whether CsrA represses translation of the *ast* transcripts. In the *A. baumannii* AB5075 Δ*csrA* background, reporter activity for *lacZ* translational fusions to *astB, astG*, and *astP* was significantly increased compared to wild-type and activity returned to wild-type levels when a wild-type copy of *csrA* was expressed *in trans* (Figures 5a-c). Importantly, ectopic expression of an RNA-binding deficient allele of CsrA (R44A) did not alter activity for any of the reporters tested (Figures 5a-c).^25^ With the above RIP-seq and EMSA data, these *lacZ-*reporter data are consistent with CsrA directly binding and acting as a translational repressor of *astGCADBE* and *astNOP* in *A. baumannii*.

**Figure 5:**
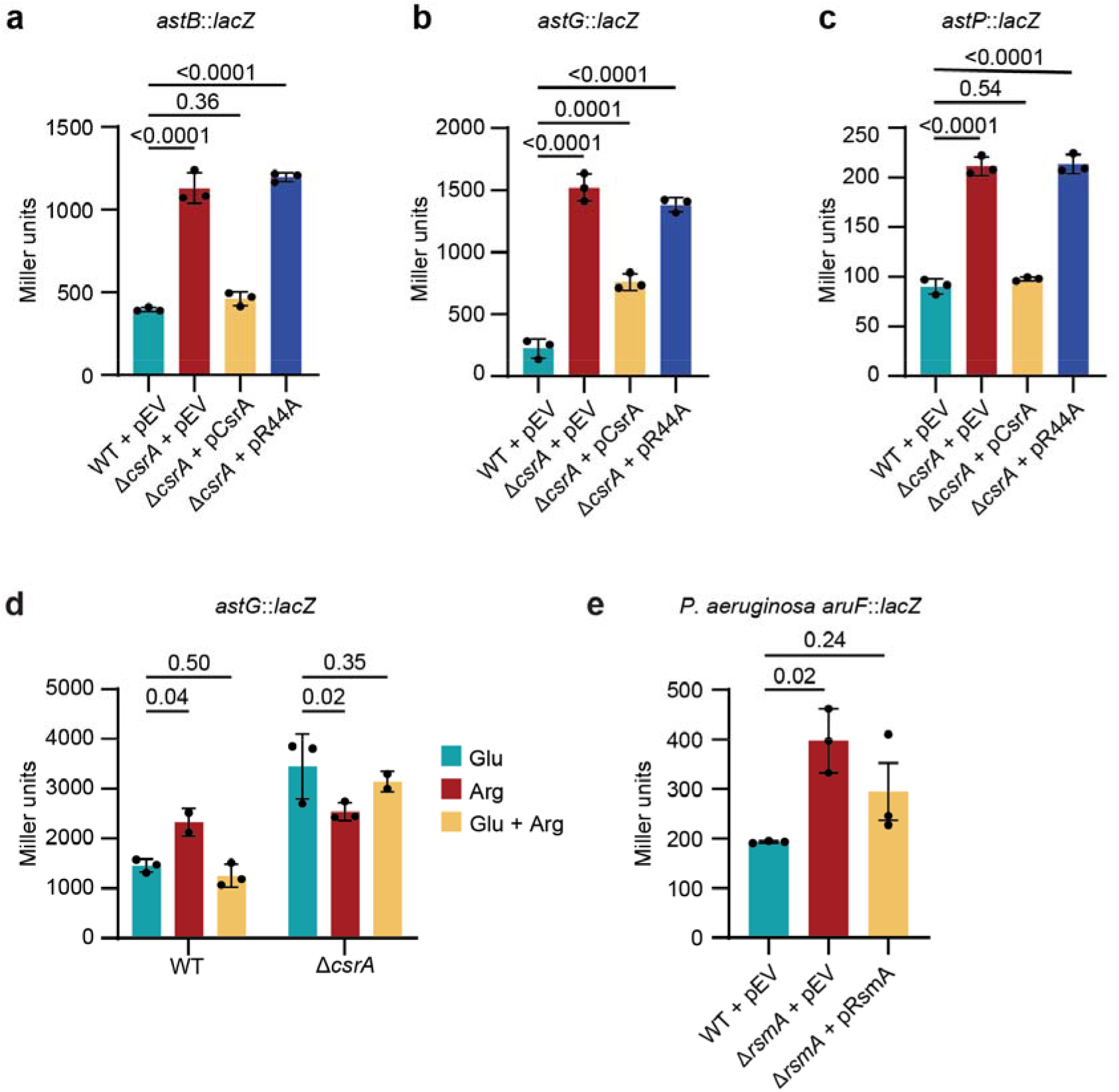
CsrA represses expression of *ast* transcripts. (a-c) Beta-galactosidase activity assays of the indicated translational reporters in AB5075 strains when grown in (a-b) M9 succinate medium or (c) M9 succinate medium plus 20 mM ornithine. P by one-way ANOVA with Dunnett’s multiple comparisons test (n = 3, mean ± SD). (d) Beta-galactosidase activity assays of *Ptac*-*astG::lacZ* translational reporters in the indicated strains when grown in M9 minimal media with 16.5 mM glutamate (Glu), 16.5 mM arginine (Arg), or 10 mM glutamate and 16.5 mM arginine (Arg + Glu) as the sole carbon sources. P by two-way ANOVA with Holm-Šídák’s multiple comparisons test (n = 2-3, mean ± SD). (e) Beta-galactosidase activity assays of *aruF::lacZ* translational reporters in the indicated *P. aeruginosa* strains when grown in M9 minimal medium with 20 mM succinate and 20 mM arginine. P by one-way ANOVA with Dunnett’s multiple comparisons test (n = 2-3, mean ± SD).

We previously reported that the *astNOP* genes are most highly expressed in succinate/ornithine medium and repressed by addition of preferred carbon sources such as glutamate.^13^ To test whether CsrA mediates similar repression of the *astGCADBE* operon, we constructed an *astG::lacZ* translational fusion driven by a heterologous inducible promoter. The *astG* reporter activity was reduced in wild-type *A. baumannii* grown in the presence of glutamate compared to growth in arginine (Figure 5d). In Δ*csrA,* however, reporter activity remained increased regardless of the presence of arginine and/or glutamate (Figure 5d). These data suggest that CsrA mediates carbon catabolite repression of the *astGCADBE* operon in the presence of glutamate, a preferred carbon source.

Previous studies characterizing global targets of CsrA regulation in other bacteria suggested that CsrA homologs repress expression of the *ast* transcripts in *E. coli*,^30^ *Salmonella enterica* subsp. Typhimurium,^31^ and *P. aeruginosa.*^32^ Intriguingly, in *P. aeruginosa*, *lacZ-*reporter fusion to *aruF* (*astA* homolog)^33^ exhibited significantly increased activity in cells lacking *rsmA* (*csrA* homolog^34^) compared to wild-type (Figure 5e); repression was restored by complementation of *rsmA* from a plasmid (Figure 5e). These data indicate that CsrA-mediated repression of the *ast* pathway is conserved beyond *Acinetobacter*.

### CsrA is required for *A. baumannii* gut colonization

We previously showed that *A. baumannii* ornithine catabolism is required for gut colonization in a mouse model but was dispensable with supplementation of preferred carbon sources glutamate or histidine.^10^ Because CsrA regulates *ast*-dependent ornithine utilization and growth in complex carbon environments, we hypothesized that CsrA would be required for gut colonization in a mouse model (Figure 6a). In the mice inoculated with *A. baumannii* 17978, there were significantly more wild-type colony forming units (CFU) than Δ*csrA* detected in the stool on days 1, 3, and 7 post-inoculation and no detectable Δ*csrA* CFU on day 9 or in the cecum or colon on day 10 (Figure 6b-c). In the mice inoculated with *A. baumannii* AB5075, there was significantly more wild-type than Δ*csrA* on days 3, 7, and 9 with no detectable Δ*csrA* CFU in the stool of any mouse on day 9 or the cecum or colon on day 10 (Figure 6d-e). These data show that *A. baumannii* requires CsrA for mouse gut colonization.

**Figure 6:**
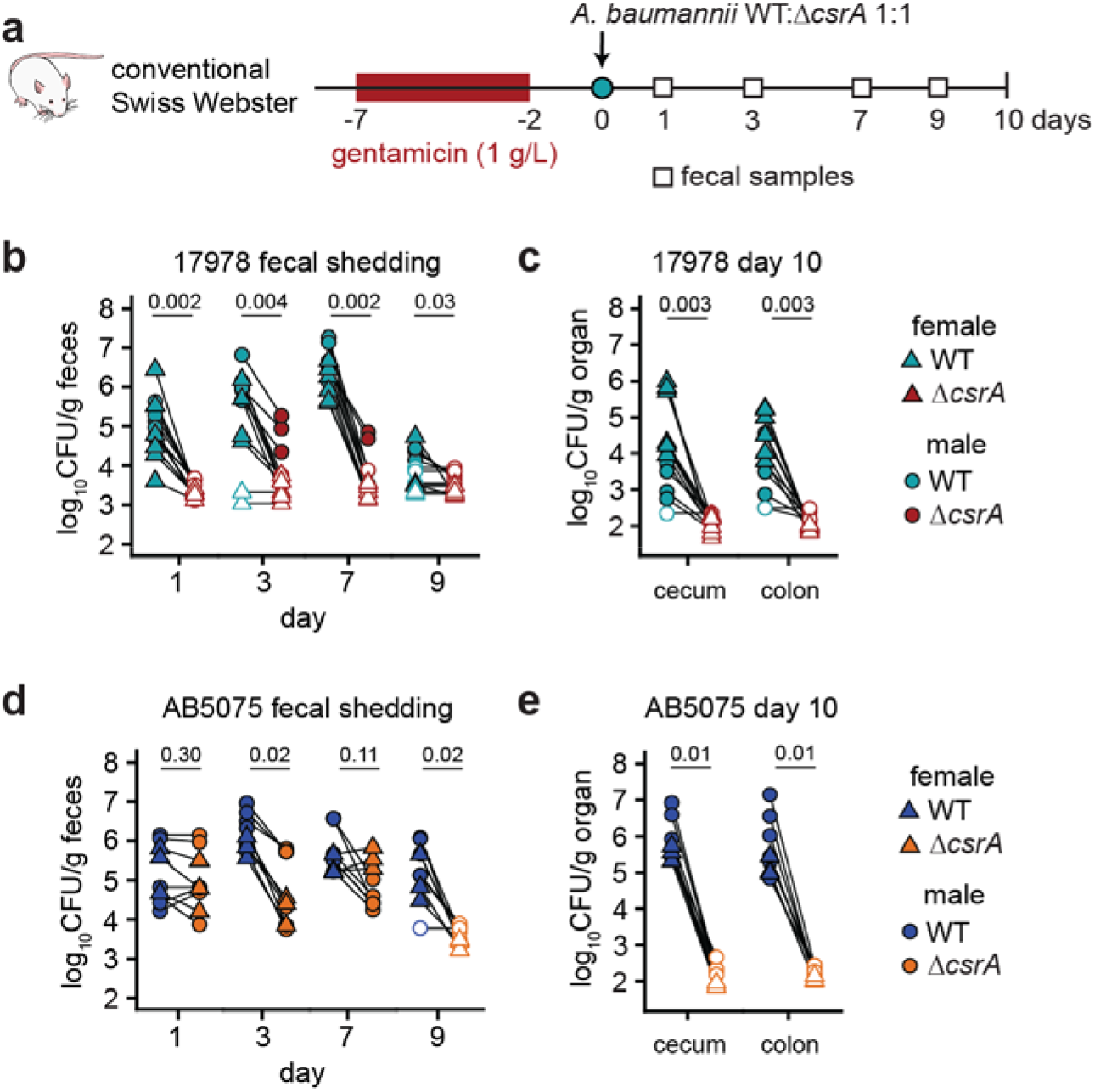
CsrA is required for *A. baumannii* gut colonization. (a) Experimental design. (b-e) Colony forming units (CFU) from fecal samples of the indicated strains at 1, 3, 7, and 9 days and organs at 10 days (17978 inoculum: n = 6 female mice, n = 6 male mice; AB5075 inoculum: n = 6 female mice, n = 3 male mice; *P* by multiple Wilcoxon tests with Holm-Sidak correction). Lines connect CFU of strains enumerated from the same mouse. Filled symbols indicate detected CFU and hollow symbols indicate undetected CFU graphed at 0.99x the limit of detection.

## Discussion

*A. baumannii* is a major multidrug-resistant pathogen for which asymptomatic colonization is an important reservoir in healthcare settings.^6^ We and others have shown that carbon metabolism and its regulation are critical drivers of *A. baumannii* gut colonization.^10,13,35,36^ Here, the data suggest that CsrA mediates *A. baumannii* carbon preference of other carbon sources over arginine, ornithine, and putrescine. CsrA associated with hundreds of transcripts during exponential growth in LB, with nearly 30% of these transcripts encoding proteins involved with metabolism. The breadth and diversity of transcripts bound by CsrA in *A. baumannii* is similar in scope with comparable experiments in CsrA from *E. coli* and RsmA from *P. aeruginosa*.^30,32^ CsrA directly bound *ast* operon RNA and was required for glutamate-mediated suppression of *astG* expression, suggesting CsrA is a carbon catabolite repressor in *A. baumannii*. The CsrA homolog RsmA also repressed *aruF* (*astA* homolog) in *P. aeruginosa*, suggesting repression of arginine/ornithine utilization is a conserved CsrA function ancestral to the Pseudomonadales family. Finally, CsrA was required for *A. baumannii* 17978 or AB5075 to persist in the mouse gut, demonstrating its importance in asymptomatic colonization.

While mechanisms for carbon preference have been well described in *E. coli*, *Bacillus subtilis*, and others, they were largely unknown in *Acinetobacter spp.*^16^ The carbon preferences reported here for *A. baumannii* are consistent with previous descriptions of “reverse CCR” in *Pseudomonas spp.* and other bacteria which use organic acids preferentially instead of sugars, which are classically preferred by *E. coli*.^16,37,38^ In *Pseudomonas spp.*, carbon preference is largely regulated by Crc,^24^ which was not required for carbon preference of glutamate over arginine or succinate in *A. baumannii* 17978. While CsrA was required for the preference of glutamate over succinate in AB5075, CsrA was not required in three other strain backgrounds. Thus, it is likely that additional regulatory mechanisms are utilized by *A. baumannii* to mediate some carbon source preferences. The extent of *A. baumannii* carbon preference mediated by CsrA across strains remains to be determined.

CsrA is a global post-transcriptional regulator known to regulate factors associated with virulence in many bacteria,^15^ however, to our knowledge CsrA has not been previously described to directly play a role in carbon preference. CsrA is associated with various phenotypes in *A. baumannii* including a phase switch associated with virulence factor expression,^26^ desiccation tolerance,^27,39^ osmotic stress tolerance,^39,40^ biofilm formation,^27,39^ amino acid metabolism,^39,40^ survival in human serum,^39^ and virulence in a *Galleria mellonella* infection model.^39^ It is likely that some of these phenotypes contribute to the requirement for CsrA in the mouse model of gut colonization. Additionally, future studies are needed to uncover the molecular details of how CsrA regulation is altered by the presence of preferred carbon sources. In *E. coli*, CsrA activity is primarily regulated by inhibitory small RNAs (sRNAs) including CsrB and CsrC.^41,42^ These sRNAs are upregulated in nutrient-poor media^43^, suggesting CsrA repression is blocked during nutrient stress to permit utilization of non-preferred carbon sources. Three sRNAs have been identified which bind CsrA in *A. baumannii*—CsrB, CsrC, and CsrD—which are regulated by the GacSA two component system.^26^ However, the conditions dictating GacSA signaling and sRNA expression in *A. baumannii* are unknown. Ultimately, CsrA is a global regulator in *A. baumannii*, and further studies are needed to better understand its functions and regulation.

In conclusion, the data herein support a model where CsrA directly binds to mRNA transcripts to mediate carbon catabolite repression of arginine and ornithine utilization in *A. baumannii*. CsrA associates with hundreds of targets and is required for *A. baumannii* gut carriage in a mouse model. Asymptomatic *A. baumannii* gut carriage presents a significant risk factor for hospitalized patients, and understanding the complex mechanisms that *A. baumannii* uses to compete with the host microbiota and persist in the gut may help develop therapeutics to prevent infections and spread.

## Acknowledgements

We thank the Palmer and Gebhardt laboratories for critical reading of the manuscript. We thank Raja Singh and Philip Rather (Emory University) for technical advice about CsrA-His purification and RNA EMSAs. We thank Elliot Friedman for assistance with amino acid analysis performed by the Microbial Culture & Metabolomics Core of the PennCHOP Microbiome Program and the UPenn Digestive and Liver Diseases (National Institutes of Health (NIH) P30DK050306). Research reported in this publication was supported by the National Institutes of Health under award numbers R35GM156848 to M.J.G and R01AI189516 and R01AI192538 to L.D.P. This work was also supported by a Pre-Doctoral Fellowship from the American Heart Association to R.M.C. (https://doi.org/10.58275/AHA.25PRE1374972.pc.gr.227132), Burroughs Wellcome Fund award number 1548433 to L.D.P., and the Michael Reese Pioneer Award to L.D.P. This manuscript is the result of funding in whole or in part by the NIH. It is subject to the NIH Public Access Policy. The content is solely the responsibility of the authors and does not necessarily represent the official views of the National Institutes of Health.

## Author Contributions

Conceptualization: R.M.C., R.L.S., M.J.G., and L.D.P.; Methodology: all authors; Investigation: R.M.C., R.L.S., and K.J.A.H.; Writing—original draft: R.M.C., R.L.S., M.J.G., and L.D.P.; Writing—review & editing: all authors; Visualization: R.M.C., R.L.S., and M.J.G.; Supervision: M.J.G. and L.D.P.; Funding acquisition: R.M.C., M.J.G., and L.D.P.

## Declaration of Interests

The authors declare no competing interests.

## Materials and Methods

### Bacterial strains and culture conditions

*A. baumannii* strains were routinely grown in Miller’s lysogeny broth (Fisher Scientific). Minimal media was carbon-free M9 minimal media with 0.1X Vishniac’s trace metals.^44,45^ Unless otherwise noted, M9 Succinate with 16.5 mM sodium succinate was used for routine culturing of Δ*csrA* mutants. Cultures were incubated with shaking at 37°C. Antibiotics were used at the following concentrations: carbenicillin 75 mg/L, kanamycin 40 mg/L, chloramphenicol 10 mg/L, hygromycin B 300 mg/L.

### Bacterial strain construction

A list of strains, plasmids, and oligonucleotides used in this study is in Table S4. Mutants in *A. baumannii* ATCC 17978^46^ were constructed by allelic exchange with sucrose counterselection using pFLP2.^47^ Oligonucleotides were purchased from Integrated DNA Technologies. The kanamycin resistance cassette was amplified from pKD4.^48^ The PCR products were cloned into the pFLP2 vector using HiFi Assembly (New England Biolabs). The pFLP2 constructs were then introduced into WT *A. baumannii* by triparental mating, and PCR-confirmed merodiploids were plated to LB agar or M9 succinate agar containing 10% sucrose to select for loss of the plasmid. Mutants were confirmed by PCR.

To construct the *A. baumannii* AB5075 Δ*csrA att::*mTn7(*hygR*) strain marked with hygromycin resistance, pJMP12089 containing Tn7(*hygR*) was transformed into *A. baumannii* AB5075 Δ*csrA* by a four parental conjugal mating strategy as described previously,^49^ using M9 succinate agar for Δ*csrA*.

Wild-type and Δ*csrA A. baumannii* 17978 was analyzed with whole genome sequencing. Short-read Illumina sequencing was performed by SeqCoast Genomics and mapped to the reference genomes using breseq v0.39.0 with bowtie2 v2.4.5.^50,51^ Reference genomes for *A. baumannii* 17978 are CP000522, CP000523, CP012004, and CP012005. Whole genome sequencing data for wild-type and Δ*csrA A. baumannii* AB5075 were reported previously.^25^ Each *csrA* deletion strain was isogenic to the wild-type strain it was compared to.

### Bacterial growth curves

*A. baumannii* strains were grown in 3 mL M9 Succinate overnight at 37°C with shaking. These cultures were diluted 1/100 into PBS, and 1 µL of this was further diluted into 100 µL M9 minimal media with the indicated carbon and nitrogen sources. Unless otherwise noted in the figure legends, the carbon source was at 16.5 mM. Bacteria were cultured in a flat-bottom 96-well plate, covered with Breathe-Easy sealing membrane, and placed in a microplate reader (Biotek Epoch or Synergy). The plates were incubated at 37°C with slow shaking, and growth was measured by optical density at 600 nm (OD_600_).

For diauxic growth curves to test carbon preference, carbon sources were assayed against glutamate, succinate, and arginine. Unless otherwise noted in the figure legends, succinate and glutamate were added to 16.5 mM of the tested carbon source, and the tested carbon source was added to 16.5 mM arginine. Lower concentrations were used when the tested carbon source was toxic at higher concentrations, as indicated.

### Experimental evolution

Twelve cultures from independent colonies of wild-type *A. baumannii* 17978 were grown overnight in 3 mL LB at 37°C with shaking. Each culture was diluted 1:2,000 daily for 10 days into M9 medium with 16.5 mM arginine as the sole carbon source. On day 10, a sample of each evolved culture was collected for gDNA extraction. Short read Illumina sequencing was performed on each sample by SeqCoast Genomics. These reads were then aligned to the wild-type reference genome using breseq v0.39.0 using the “-p” option to predict polymorphic mutations.^50,51^

The breseq output was manually transferred into a dataframe and analyzed in RStudio v2025.09.2 using R v4.5.2 using code available below. In short, each mutation or new predicted junction was paired with the locus tag where it was detected. Intergenic mutations were assigned to both the locus upstream and downstream from the detected mutation. If multiple mutations were found in a locus tag in the same population, the mutation with the highest prevalence was selected. The data were filtered for locus tags that had mutations detected in at least 6 independent populations and had a mutation with a prevalence of at least 25% in at least one population. Loci known to be part of mobile genetic elements were manually relabeled in Figure 2C for clarity (*i.e.* ISAba*1*).

### Amino acid metabolomics

Four independent cultures of *A. baumannii* 17978 WT and Δ*csrA* were grown overnight in 3 mL M9 Succinate. The cultures were centrifuged at 4,000 x *g* and resuspended in equal volume PBS twice. The cultures were then diluted 1:50 into carbon-free nitrogen-free M9 with 18.6 mM ammonium chloride and 5 mM each glutamate, arginine, and ornithine as the sole carbon sources. These cultures were incubated at 37°C with shaking and 100 uL aliquots were taken for OD_600_ measurement every 30 min for 8 h and one sample at 28 h. Additional supernatant was collected at 0, 2.5, 5, 8, and 28 h, sterile filtered, and frozen at -80°C. Amino acid metabolomics was performed on these samples by the Microbial Culture & Metabolomics Core of the PennCHOP Microbiome Program and the UPenn Digestive and Liver Diseases. Amino acids were derivatized using the Waters AccQ-Tag Ultra Amino Acid Derivatization Kit (Waters Corporation) and analyzed using the UPLC AAA H-Class Application Kit (Waters Corporation, Milford, MA) according to manufacturer’s instructions. Amino acids were quantified using a Waters Acquity UPLC System with an AccQ-Tag Ultra C18 1.7 μm, 2.1 × 100 mm column and a photodiode detector array with a standard curve. Quality control checks (blanks and standards) were run every eight samples.

### RNA-immunoprecipitation sequencing (RIP-seq)

Samples for RNA Immunoprecipitation (RIP) were collected as previously described.^25,52,53^ Briefly, 80 mL of triplicate mid-log cultures (OD_600_= 0.5 or 40 OD units) of wild-type AB5075 and a strain encoding a Vesicular stomatitis virus (VSV) G protein epitope from the C-terminus of the *csrA* allele (CsrA-V) grown in LB were collected by centrifugation. Cell pellets were washed with PBS and exposed to ultraviolet light to induce crosslinking. Cross-linked cells were lysed by bead beating and an aliquot of sample was taken for total RNA sequencing prior to immunoprecipitation (IP). IP was performed on cells lysates with murine monoclonal anti-VSV-G antibodies (Sigma Aldrich #SAB4200695). Following IP, proteins were digested with Proteinase K (ThermoFisher). IP and Total RNA was then extracted with TriReagent LS according to the manufacturer’s recommendations and GlycoBlue (Ambion) was added to aid in RNA precipitation. Samples were DNAse treated with Invitrogen DNAse (RNAse free). Library preparation was performed using Illumina’s Stranded Total RNA Prep Ligation with 10bp unique dual indices (UDI). Ribosomal RNAs (rRNAs) were not depleted during sequencing library construction. Sequencing was performed on a NovaSeq X Plus, producing paired end 150bp reads. Demultiplexing, quality control, and adapter trimming was performed with bcl-convert (v4.2.4).

RNA reads from RIP-Seq samples were mapped to the AB5075 genome using bowtie2 (version 2.51.1) using the following settings: -N 1 -L 28 -i S,1,2.00 -D 20 --non-deterministic --dovetail --no- mixed --no-discordant --no-unal. Mapped reads were converted to bam format and reads aligned to the rRNA regions were removed using samtools. Filtered reads were further analyzed for enrichment using the ‘callpeak’ command in MACS3^54^ with keep duplicates set to auto, an effective genome size of 3.9e6, and a false discovery (q-value) cutoff of 0.01. Following peak calling, custom scripts were used to compile the peak number, enrichment factor, genome coordinates for max peak height, peak start and end coordinates, peak size, gene locus, strand, gene name, and gene product. A Circos^55^ plot was used to visualize reads across the genome and IGV^56^ was used to visualize reads and enrichment regions.

### Total RNA Sequencing

Total RNA was extracted from WT and CsrA-V samples collected for RIP-Seq as described above. Sequencing library preparation was performed as described above with the addition of an rRNA removal step with the Ribo-Zero Plus kit. Reads were aligned to the AB5075 genome using Bowtie2 using the following settings: -N 1 -L 28 -i S,1,2.00 -D 20 --non-deterministic --dovetail --no-mixed --no-discordant --no-unal. The alignment files were normalized for read depth and converted to bigwig format using custom scripts to facilitate visualization in IGV.

### CsrA-His purification

The wild-type *csrA* sequence was cloned into the pET21a vector (NovaGen) by HiFi Assembly (New England Biolabs). Sequence-verified pET21a-CsrA-His was transformed into Arctic Express (DE3) RIL competent *Escherichia coli* (Agilent). This strain was grown in 10 mL LB with 20 mg/L gentamicin and 50 mg/L carbenicillin and incubated overnight at 37°C with shaking. The culture was then diluted 1:100 into 1 L LB containing 50 mg/L carbenicillin and incubated at 37°C with shaking. At OD_600_ = 0.6, IPTG was added to 1 mM. After 3 h incubation at 37°C with shaking, the cells were centrifuged at 4,000 x *g* and the supernatant was discarded. The pellet was then frozen at -80°C until the next day. Pelleted cells were thawed and lysed using Bacterial Protein Extraction Reagent Complete (B-PER, Thermo Scientific) per the manufacturer’s instructions using 10 mL/gram cell pellet.

Next, the soluble lysate was mixed 1:1 with lysis buffer (50 mM NaH_2_PO_4_, 300 mM NaCl, 10 mM imidazole, pH 8.0), mixed with 3 mL equilibrated Ni-NTA resin (Qiagen), rocked at 4°C for 1 h, and loaded onto 2 gravity columns with a 1.5 mL bed volume. Each column was then washed twice with 7.5 mL wash buffer (50 mM NaH2PO4, 300 mM NaCl, 20 mM imidazole, pH 8.0). Finally, each column was eluted four times with 1.5 mL elution buffer (50 mM NaH2PO4, 300 mM NaCl, 250 mM imidazole, pH 8.0). Fractions were analyzed by SDS-PAGE, and the fractions containing purified CsrA-His were pooled. Finally, the pooled fractions were desalted using PD-10 columns (Cytiva) per the manufacturer’s instructions and eluted in a total of 3.5 mL storage buffer (10 mM HEPES pH 7.3, 20 mM KCl, 10% v/v glycerol). Protein concentration was quantified using Pierce 660nm Protein Assay with bovine serum albumin standards as reference (Thermo Scientific).

### RNA electromobility shift assays (EMSA)

RNA probes were generated by PCR to amplify the target regions with a T7 promoter at the 5’ end. RNA was then transcribed using the HiScribe T7 High Yield RNA Synthesis Kit (New England Biolabs) and purified using the Monarch Spin RNA Cleanup Kit (New England Biolabs). RNA was eluted in water and concentrations were determined with a Nanodrop spectrophotometer (Thermo Scientific).

RNA EMSA was performed as previously described using the LightShift Chemiluminescent RNA EMSA kit (Thermo Scientific).^26^ 20 ng RNA was used in each reaction along with the indicated concentration of CsrA-His. Each reaction was incubated for 30 min at room temperature and then separated on a 5% Mini-Protean TBE gel (Bio-Rad) with 0.5X TBE running buffer at 100 V for 100 min. The gels were then stained with Sybr Gold (Thermo Scientific) for 30 min and imaged with a Chemi-doc MP imager (Bio-Rad).

### Reporter assays

β-galactosidase assays were conducted as previously described.^25^ Overnight cultures of the indicated reporter were back diluted 1:50 into fresh growth media and grown to early exponential phase (OD 0.1-0.2). Bacterial cells were then permeabilized with chloroform and 0.1% SDS. β-galactosidase activity was assayed using 2-nitro-phenyl-β-D-galactopyranoside (ONPG). Assays were repeated independently at least three times.

### Mouse experiments

All animal experiments were performed in compliance with NIH guidelines, the Animal Welfare Act, and US federal law. All animal experiments were approved by the Institutional Animal Care and Use Committee in protocol 23-119 at the Biological Resources Laboratory of the University of Illinois Chicago. All mice were euthanized by methods consistent with American Veterinary Medical Association (AVMA) guidelines.

Mouse gut colonization was performed as previously described.^10^ Six-week-old female Swiss-Webster mice from Inotiv (strain Hsd:ND4) or male Swiss-Webster mice from Charles River Laboratories (strain Crl:CFW(SW)) were acclimated to the facility for one week. Mice were fed Teklad Irradiated LM-485 mouse diet and water *ad libitum*. The mice were housed with 14 h:10 h light:dark cycles, 70–76°F and 30%–70% humidity. After one week, gentamicin was administered at 1 g/L in the drinking water *ad libitum* for 5 days (days -7 to -2). The mice were allowed to recover with normal drinking water for two days before inoculation (days -2 to 0).

*A. baumannii* was prepared for inoculation by growing overnight cultures in 10 mL M9 Succinate. The cultures were pelleted at 4,000 x *g* for 10 minutes and resuspended in equal volumes of PBS twice. The culture was then pelleted and resuspended in PBS to approximately 3 x 10^10^ CFU/mL. Mice were then inoculated by orogastric gavage with 100 µL of a 1:1 mixture of either *A. baumannii* 17978 WT *att*::mTn*7*(Carb^R^) and *A. baumannii* 17978 Δ*csrA*::Kn or *A. baumannii* AB5075 WT and *A. baumannii* AB5075 Δ*csrA att*::Tn*7*(Hyg^R^). Resuspended cultures were serially diluted and plated to quantify each inoculum.

Selective media for plating and quantifying each strain was as follows: *A. baumannii* 17978 WT *att*::mTn*7*(Carb^R^) on LB agar with chloramphenicol 5 mg/L and carbenicillin 50 mg/L; *A. baumannii* 17978 Δ*csrA*::Kn on M9 Succinate agar with chloramphenicol 5 mg/L and kanamycin 40 mg/L; *A. baumannii* AB5075 WT on LB agar with chloramphenicol 10 mg/L; and *A. baumannii* AB5075 Δ*csrA att*::Tn*7*(HygR) on M9 Succinate agar with hygromycin B 300 mg/L. Fecal samples were collected from each mouse prior to inoculation, homogenized in PBS, and plated on selective media to confirm that there were no detectable CFU.

Fecal pellets were weighed and homogenized manually in 1 mL PBS with sterile wooden sticks. On day 10, mice were humanely euthanized by carbon dioxide. Organ samples were collected, weighed, and homogenized in 1 mL PBS in 1.5 mL Bullet Blender navy tubes (NextAdvance). Sample weights were used to calculate average limits of detection (LOD) for each sample type. Fecal and organ samples were serially diluted for CFU counting on selective media.

### Data reporting, statistical analysis, and figure preparation

Each measurement was taken from a distinct biological sample (for example, an individual mouse or bacterial culture from a single colony). For CFU enumeration, samples below the limit of detection are analyzed and graphed at 0.99X the limit of detection. Data processing, statistical analyses, and graphing were performed with Microsoft Excel version 2602, GraphPad Prism version 11.0.0, or R version 4.5.2 with RStudio version 2025.09.2. Statistical tests are listed in each figure legend. Figures were prepared using Adobe Illustrator version 30.2.1.

## Data availability

Whole genome sequencing data for experimental evolution populations and for *A. baumannii* 17978 Δ*csrA* can be found at NCBI sequence read archive (SRA) accession number PRJNA1548715. Raw sequencing reads for the CsrA-V RIP-seq and corresponding total RNA-seq have been deposited at the NCBI GEO database and will be available upon formal publication.

## Code availability

The R code used for the experimental evolution analysis of breseq outputs can be found at https://github.com/RMasonClark/ExptEvoAnalysis. The scripts used for RIP-seq peak calling and RNA-seq visualization can be found at: https://github.com/mjgeby/A.-baumannii-RIP-seq-peaks.

## Notes

### Competing Interest Statement

The authors have declared no competing interest.

