## Supplemental Figures for "A global post-transcriptional regulator directs *Acinetobacter baumannii* carbon preference and gut colonization"

**Supplementary Tables:**

**Table S1:** Growth curve data for *A. baumannii* 17978 carbon preferences shown in Figure 1a

**Table S2:** Population sequencing identified mutations and rearrangements in experimental evolution

**Table S3:** RIP-seq targets of CsrA-V in AB5075

**Table S4:** Strains, plasmids, and oligonucleotides

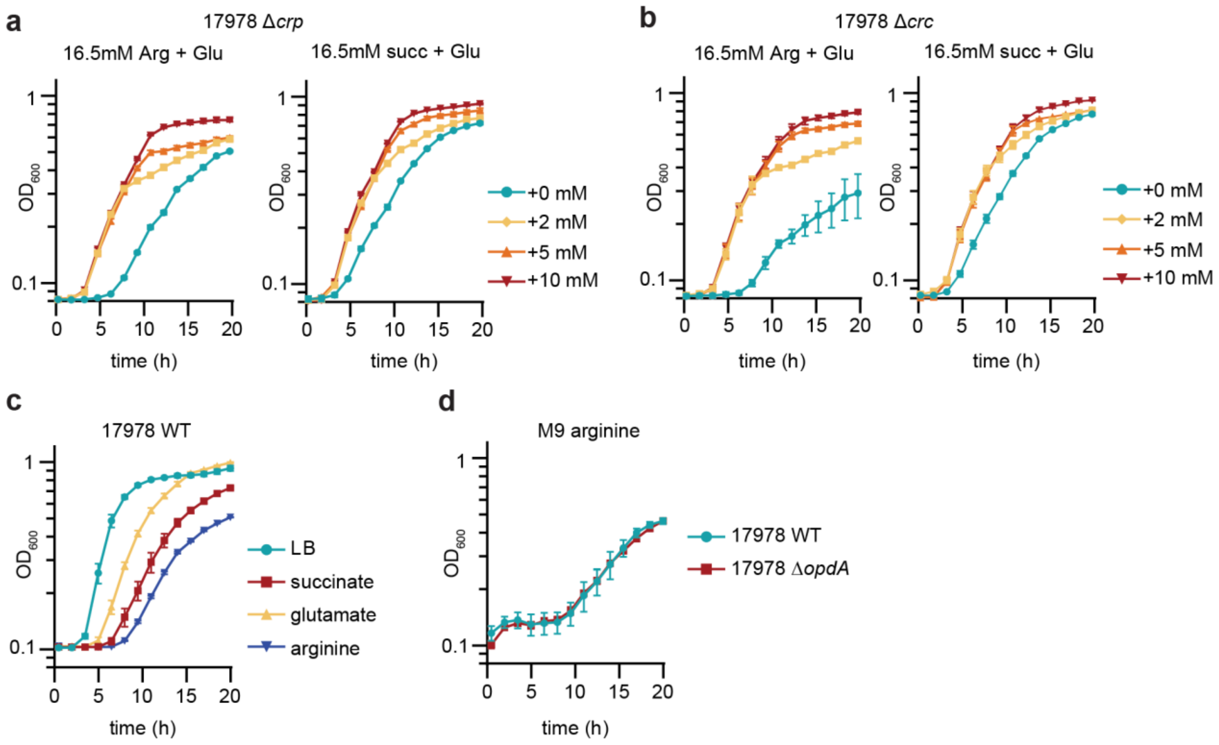


**Figure S1: Neither *crp* nor *crc* are required for *A. baumannii* carbon preference, and *opdA* does not affect growth on arginine minimal media.**

(a-b) Diauxic growth curves of the indicated *A. baumannii* strain in M9 media with either 16.5 mM arginine or 16.5 mM succinate and the indicated concentration of glutamate as sole carbon source(s). Growth was monitored by optical density at 600 nm ($OD_{600}$, $n$ = 3, mean ± SEM, experiments were repeated at least twice with similar results).

(c) Growth curves of wild-type *A. baumannii* 17978 in either LB or M9 media with the indicated compound added at 16.5mM as the sole carbon and nitrogen source. Growth was monitored by optical density at 600 nm ($OD_{600}$, $n$ = 3, mean ± SEM, experiments were repeated at least twice with similar results).

(d) Growth curves of the indicated strain of *A. baumannii* in M9 arginine media. Growth was monitored by optical density at 600 nm ($OD_{600}$, $n$ = 3, mean ± SEM, experiments were repeated at least twice with similar results).

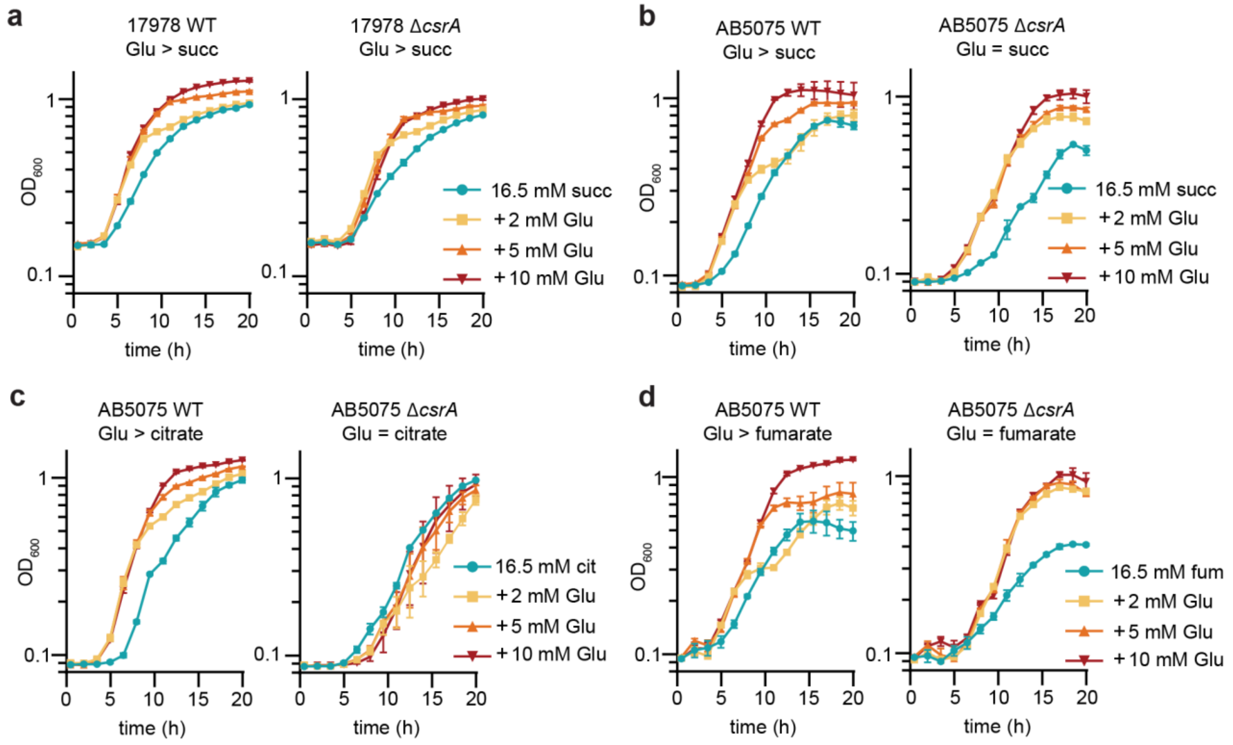


**Figure S2: CsrA is required for additional carbon preferences in AB5075.**

(a-d) Diauxic growth curves of the indicated *A. baumannii* strains in M9 media with the indicated concentrations of carbon source(s). Growth was monitored by optical density at 600 nm ($OD_{600}$, $n$ = 3, mean ± SEM, experiments were repeated at least twice with similar results).

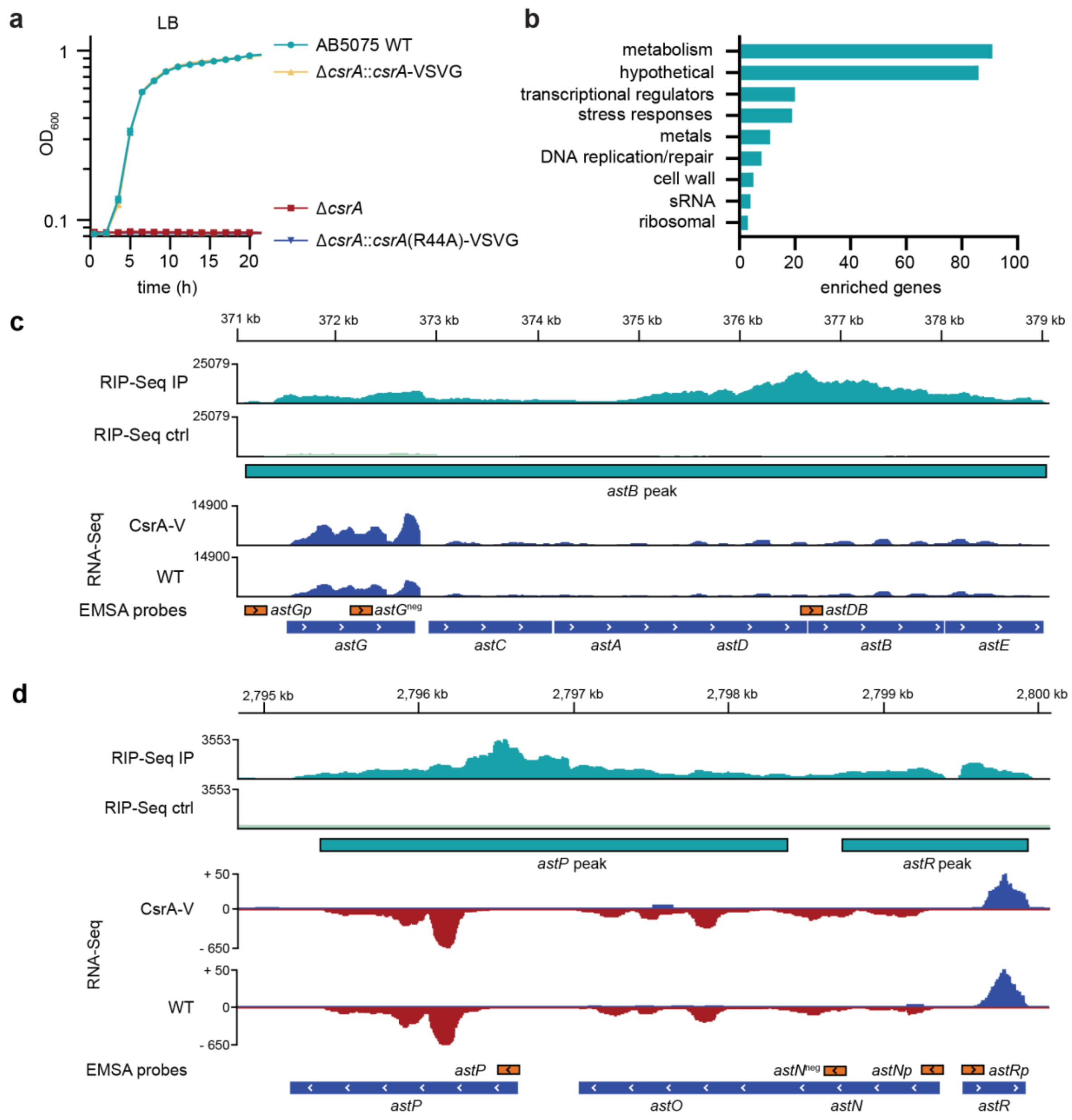


**Figure S3: RIP-seq identified both *ast* transcripts as CsrA binding targets**

(a) Growth curves of the indicated strains of *A. baumannii* AB5075 in LB media. Growth was monitored by optical density at 600 nm ($OD_{600}$, $n$ = 3, mean ± SEM, experiments were repeated at least twice with similar results).

(b) Functional analysis of RNA targets enriched ≥ 1.5-fold in the CsrA RIP-seq experiment (n = 289 targets).

(c-d) RIPseq traces showing enriched reads overlayed on the *ast* operons as shown in Figure 4c-d. Additionally shown are the corresponding control RIP-Seq read density trace and total RNA sequencing reads. RNA probes used in RNA EMSAs and the *ast* operon structure are shown below the RNA sequencing traces.

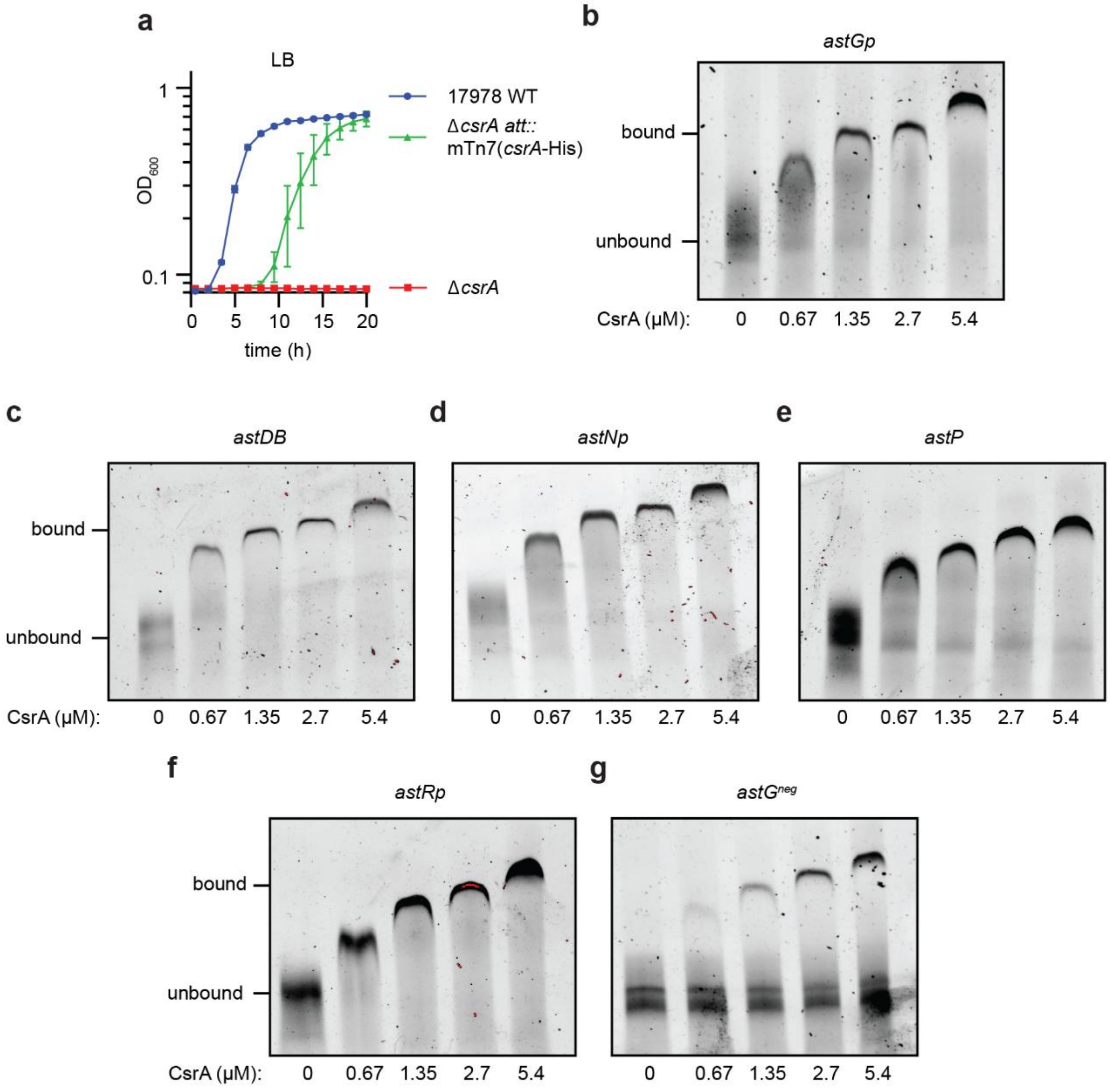


**Figure S4: CsrA binds multiple loci in both *ast* RNA transcripts**

(a) Growth curves of the indicated strains of *A. baumannii* 17978 in LB media. Growth was monitored by optical density at 600 nm ($OD_{600}$, $n$ = 3, mean ± SEM, experiments were repeated at least twice with similar results).

(b-g) EMSA analysis of purified CsrA-His binding to RNA probes corresponding to the *ast* transcripts. The final concentration of CsrA-His is shown below each lane.
